# mPies: a novel metaproteomics tool for the creation of relevant protein databases and automatized protein annotation

**DOI:** 10.1101/690131

**Authors:** Johannes Werner, Augustin Geron, Jules Kerssemakers, Sabine Matallana-Surget

**Affiliations:** Department of Biological Oceanography, Leibniz Institute for Baltic Sea Research, Rostock, 18119, Germany; Division of Biological and Environmental Sciences, Faculty of Natural Sciences, University of Stirling, Stirling, FK9 4LA, UK; Proteomic and Microbiology Department, University of Mons, Mons, 7000, Belgium; Omics IT and Data Management, German Cancer Research Center, Heidelberg, 69120, Germany

**Keywords:** Bioinformatics, Metaproteomics, Microbial ecology, Protein annotation, Protein search database

## Abstract

Metaproteomics allows to decipher the structure and functionality of microbial communities. Despite its rapid development, crucial steps such as the creation of standardized protein search databases and reliable protein annotation remain challenging. To overcome those critical steps, we developed a new program named mPies (**m**eta**P**roteomics **i**n **e**nvironmental sciences). mPies allows the creation of protein databases derived from assembled or unassembled metagenomes, and/or public repositories based on taxon IDs, gene or protein names. For the first time, mPies facilitates the automatization of reliable taxonomic and functional consensus annotations at the protein group level, minimizing the well-known protein inference issue which is commonly encountered in metaproteomics. mPies’ workflow is highly customizable with regards to input data, workflow steps, and parameter adjustment. mPies is implemented in Python 3/Snakemake and freely available on GitHub: https://github.com/johanneswerner/mPies/.

## Implementation

### Introduction

Metaproteomics is a valuable method to link the taxonomic diversity and functions of microbial communities [1]. However, the use of metaproteomics still faces methodological challenges and lacks of standardisation [2]. The creation of relevant protein search database and protein annotation remain hampered by the inherent complexity of microbial communities [3].

Protein search databases can be created based on reads or contigs derived from metagenomic and/or metatranscriptomic data [4, 5]. Public repositories such as Ensembl [6], NCBI [7] or UniProtKB [8] can also be used as search databases but it is necessary to apply relevant filters (e.g. based on the habitat or the taxonomic composition) in order to decrease computation time and false discovery rate [4]. Until now, no tool exists that either creates taxonomic or functional subsets of public repositories or combines different protein databases in order to optimize the total number of identified proteins.

The so-called protein inference issue occurs when the same peptide sequence is found in multiple proteins, thus leading to inaccurate taxonomic and functional interpretation [9]. To address this issue, protein identification software tools such as ProteinPilot (Pro Group algorithm) [10], Prophane [11] or MetaProteomeAnalyzer [12] perform automatic grouping of homologous protein sequences. Interpreting protein groups can be challenging especially in complex microbial community where redundant proteins can be found in a broad taxonomic range. A well-known strategy to deal with homologous protein sequences is to calculate the lowest common ancestor (LCA). For instance, MEGAN performs taxonomic binning by assigning sequences on the nodes of the NCBI taxonomy and calculates the LCA on the best alignment hit [13]. However, another crucial challenge related to protein annotation still remains: protein sequences annotation often relies/or uses on the alignment programs automatically retrieving the first hit only [14]. The reliability of this approach is hampered by the existence of taxonomic and functional discrepancies among the top alignment results with very low e-values [5].

Here, we present mPies, a highly customizable workflow that allows designing suitable protein search databases as well as facilitating biological interpretation of identified proteins. mPies can create non-redundant databases derived from metagenomic data and public protein repositories subsets based on lists of taxon, gene and protein IDs. mPies automatically computes consensus annotation at the protein group level by taking into account multiple alignment hits for reliable taxonomic and functional inference.

### Workflow design

mPies provides a standardized and flexible workflow for both the creation of protein search database and subsequent protein annotation (Figure 1). It is written in Python, uses the workflow management system Snakemake [15] and relies on the Bioconda distribution [16] to ensure reproducibility.

**Figure 1:**
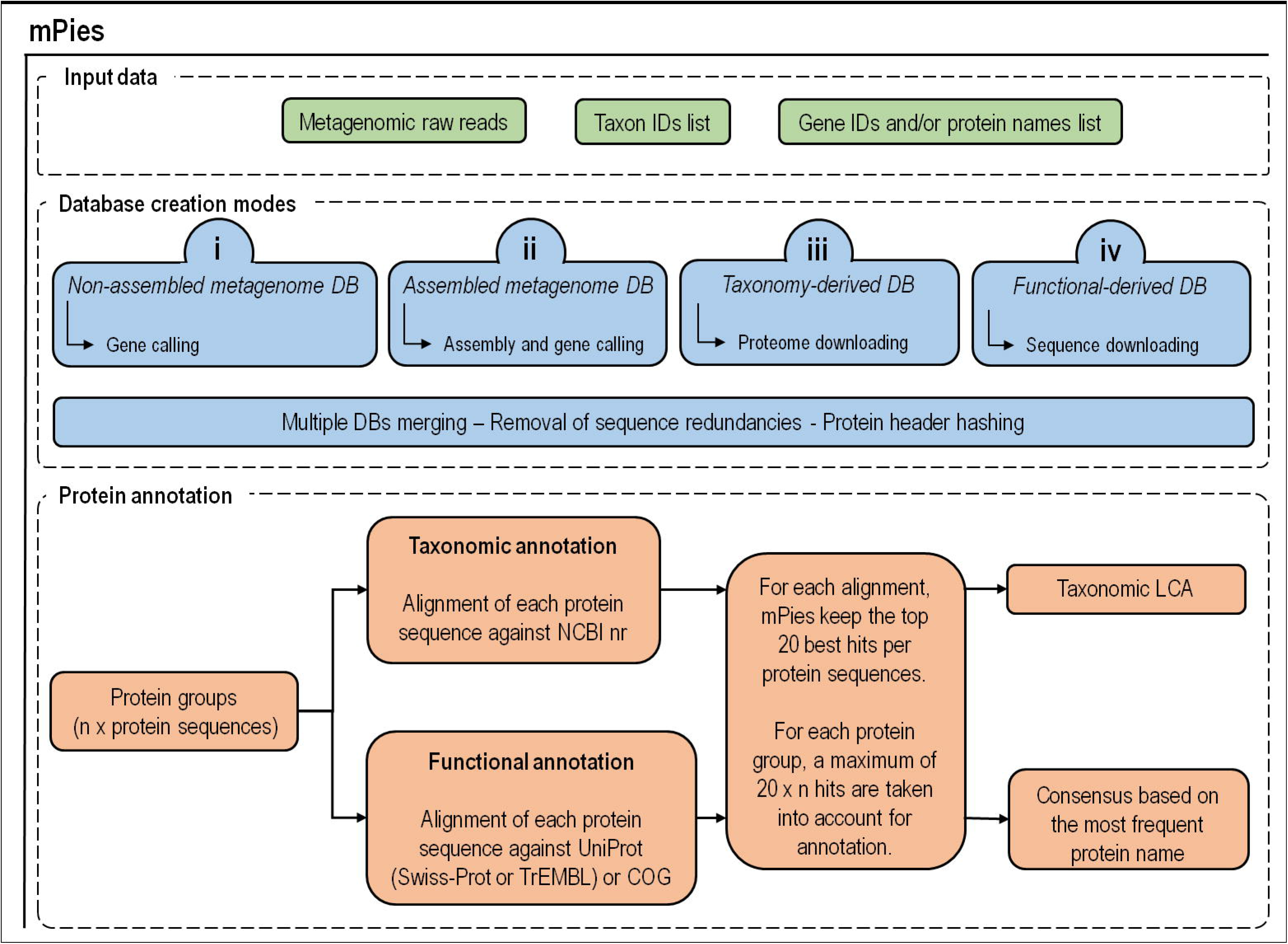
Workflow of mPies.

mPies can run in up to four different modes to create protein search databases (DBs): nonassembled metagenome-derived DB, assembled metagenome-derived DB, taxonomy-derived DB, and functional-derived DB.

#### Mode (i): Non-assembled metagenome-derived DB

In mode (i), mPies trims metagenomic raw reads (fastq files) with Trimmomatic [17], and predicts partial genes with FragGeneScan [18] which are built into the protein DB.

#### Mode (ii): Assembled metagenome-derived DB

In mode (ii), trimmed metagenomic reads are assembled either with MEGAHIT [19] or metaSPAdes [20]. The genes are subsequently called with Prodigal [21]. The utilization of Snakemake allows easy adjustment of the assembly and gene calling parameters.

#### Mode (iii): Taxonomy-derived DB

In mode (iii), mPies extracts the taxonomic information derived from the metagenomic raw data and downloads the corresponding proteomes from UniProt. To do so, mPies uses SingleM [22] to predict OTUs from the metagenomic reads. Subsequently, a non-redundant list of taxon IDs corresponding to the taxonomic diversity of the observed habitat is generated. Finally, mPies retrieves all available proteomes for each taxon ID from UniProt. It is noteworthy that the taxonomy-derived DB can be generated from 16S amplicon data or a user-defined list.

#### Mode (iv): Functional-derived DB

Mode (iv) is a variation of mode (iii) which allows to create DBs that target specific functional processes (e.g. carbon fixation or sulphur cycle) instead of downloading entire proteomes for taxonomic ranks. For that purpose, mPies requires a list of gene or protein names as input and downloads all the corresponding protein sequences from UniProt. Taxonomic restriction can be defined (e.g. *Proteobacteria-related* sequences only) for highly specific DB creation.

### Post-processing

If more than one mode was selected for protein DB generation, all proteins are merged into one combined protein search DB. Duplicated protein sequences (default: sequence similarity 100%) are removed with CD-HIT [23]. All protein headers are hashed (default: MD5) to obtain uniform headers and to reduce the file size for the final protein search database in order to keep the memory requirements of downstream analysis low.

### Protein annotation

mPies facilitates taxonomic and functional consensus annotation at protein level. After protein identification, each protein is aligned with Diamond [24] against NCBI-nr [25] for the taxonomic annotation. For the functional prediction, proteins are aligned against UniProt (Swiss-Prot or TrEMBL) [8] and COG [26]. The top 20 alignment hits (bitscore ≥ 80) are automatically retrieved for consensus taxonomic and functional annotation, for which the detailed strategies are provided below.

The taxonomic consensus annotation uses the alignment hits against NCBI-nr and applies the LCA algorithm to retrieve a taxonomic annotation for each protein group (protein grouping comprises the assignment of multiple peptides to the same protein and is facilitated by proteomics software) as described by Huson *et al.* [13]. For the functional consensus, the alignment hits against UniProt and/or COG are used to extract the most frequent functional annotation per protein group. This is the first time that a metaproteomics tool includes this critical step, as previously only the first alignment hit was kept. This level of accuracy provided by mPies delivers a more reliable taxonomic and functional annotation.

### Conclusions

The field of metaproteomics has rapidly expanded in recent years and has led to valuable insights in the understanding of microbial community structure and functioning. In order to cope with metaproteomic limitations, new tools development and workflow standardization are of urgent needs. With regards to the diversity of the technical approaches found in the literature which are responsible for methodological inconsistencies and interpretation biases across metaproteomic studies, we developed the open-source program mPies. It proposes a standardized and reproducible workflow that allows customized protein search DB creation and reliable taxonomic and functional protein annotations. mPies facilitates biological interpretation of metaproteomics data and allows unravelling microbial community complexity.

## Availability and requirements

**Project name:** mPies

**Project homepage:** https://github.com/johanneswerner/mPies/

**Operating system:** Linux

**Programming language:** Python 3

**Other requirements:** Snakemake, bioconda

**License:** GNU GPL

**Any restrictions to use by non-academics:** none

## Declarations

### Ethics approval and consent to participate

Not applicable

### Consent for publication

Not applicable

### Availability of data and material

Not applicable

### Authors ‘ contributions

JW, SMS, and AG designed mPies. JW developed and implemented mPies, JK contributed valuable discussions with regards to the software design of mPies and performed code review. JW, AG and SMS wrote substantial parts of the manuscript, all authors performed proofreading and approved the final version of the manuscript.

## Acknowledgements

This work was supported by the Royal Society funded Research Grant [RG160594]. The authors acknowledge the use of de.NBI cloud and the support by the High Performance and Cloud Computing Group at the Zentrum für Datenverarbeitung of the University of Tübingen and the Federal Ministry of Education and Research (BMBF) through grant no 031 A535A. JW and JK want to personally acknowledge Manuel Prinz and Katrin Leinweber (Technische Informationsbibliothek, TIB.eu) for code review, critical thoughts, and software publication advice.

